# Perceptual and neural biomarkers of distraction from an external sound source are not associated with tinnitus severity

**DOI:** 10.64898/2026.01.27.702073

**Authors:** David O. Sorensen, Jenna A. Sugai, Stéphane Maison, Kenneth E. Hancock, Daniel B. Polley

## Abstract

Individuals with chronic tinnitus perceive a phantom sound that imposes either a bothersome and irrepressible distraction throughout waking hours or a relatively mild nuisance that often fades into subliminal awareness. The difference in tinnitus salience may reflect a general difference in inhibitory control over any distracting sound, whether externally or internally generated. To test this hypothesis, we investigated neural and behavioral signatures of external auditory distraction suppression in participants with chronic tinnitus that had mild or bothersome tinnitus but were otherwise matched for age and hearing loss. Participants in both groups underwent behavioral and EEG testing that asked them to report on a target stream of amplitude modulated tones that switched from a random arrangement to a repeating sequence. Using additional sounds that imposed varying levels of distraction, we documented neural and perceptual suppression of auditory distractors. Behaviorally, participants with mild versus bothersome tinnitus showed comparable reductions in accuracy in the presence of varying distractor loads. Neural synchronization to the target stimulus change rate provided a useful proxy for distraction effects but did not differ between tinnitus groups. Likewise, no group differences were observed in the neural synchronization to modulation rates of the target or distractor stimuli. Our results build on work showing that individuals with tinnitus perform as well as individuals with normal hearing on listening tasks in noisy environments and expand this observation into the neural representation of sounds. Suppression of the internally generated phantom percept does not appear to be linked to general deficits in suppressing distractors.

## Introduction

Individuals with tinnitus perceive a constant, indefatigable phantom sound. For some of these individuals, the phantom sound often recedes from conscious awareness unless it is attended to or is heighted during moments of elevated stress. For others, though, the sound is more constant, interfering with sleep, focused attention, and general mental wellness (Newman et al., 1996). Understanding the differences between the more benign and globally bothersome presentations of tinnitus may be informative in designing and evaluating tinnitus treatments.

Many different models for tinnitus have been proposed (Clarke et al., 2023; Ghodratitoostani et al., 2016; Hallam et al., 1984; Jastreboff, 1990; McKenna et al., 2014). Among these various models, common components include a sensory insult that results in the phantom percept; a negative affective response that develops to the phanom percept; and cognitive or attentional changes that lead to poor suppression of the phantom percept. Efforts have been undertaken to explore how these components manifest both behaviorally and neurophysiologically.

The leading model for the biological basis of the tinnitus percept posits that a disruption in the normal balance of excitation and inhibition within auditory processing centers causes excess neural gain and hyper-synchrony that manifests in the perception of a sound without a physical source (Eggermont & Roberts, 2004; Hayes et al., 2021; Yang et al., 2011). In humans, much of the work focused on the phantom percept has involved exploration of the psychophysical properties of the tinnitus percept. Methods have been developed for participants to match their phantom percept across a number of features, including loudness and frequency (Henry & Meikle, 2000). Alternatively, the loudness of tinnitus can be assessed by adjusting the level of an external noise stimulus until it masks the tinnitus percept (Feldmann, 1971; Zagólski & Strek, 2014). These attempts to directly measure the tinnitus percept have shown only weak associations, at best, with tinnitus distress (Martines et al., 2010; Smalt et al., 2022; Wallhäusser-Franke et al., 2012). An attempt to directly measure central gain in humans did differentiate individuals with tinnitus and/or hyperacusis from a matched neurotypical sample but showed minimal association with individual differences in tinnitus distress (Smith et al., 2025).

More recent studies seeking to find objective markers of tinnitus have focused more on affective and cognitive factors. For example, pupil and facial motion markers related to emotionally evocative sounds have been shown to predict self-reported tinnitus and hyperacusis severity (Smith et al., 2025). This builds on models of tinnitus that emphasize the role of the limbic system in tinnitus distress (Rauschecker et al., 2010). Other work has shown that auditory evoked potentials are differentially modulated by attention in participants with tinnitus compared to those without tinnitus (Delb et al., 2008; Jacobson et al., 1996; Richardson et al., 2024). Together, these studies show the promise of focusing on higher-order effects of tinnitus in the search for biomarkers that can inform treatment efficacy.

More generally, studies looking at how tinnitus presence affects executive function have consistently found deficits in different aspects of executive function and attention (Andersson et al., 2000; Hallam et al., 2004; Heeren et al., 2014; Mohamad et al., 2016; Stevens et al., 2007). Many of these studies have focused on domain general cognitive and attentional effects, although it’s possible there may be modality-specific effects (Husain et al., 2015). In particular, as individuals with tinnitus seemingly vary in their ability to suppress the tinnitus percept, this may generalize to the suppression of externally presented distractors (Cuny et al., 2004; Jacobson et al., 1996; Lima et al., 2020). There is some evidence, for example, that suggests that individuals with tinnitus perform slightly worse in speech-in-noise tasks compared to matched cohorts without tinnitus (Madhukesh et al., 2024). Speech-in-noise tests have relatively high ecological validity but also involve several confounding factors: Language networks and other cognitive resources are brought to bear by speech tasks, and most maskers provide some level of both energetic and informational masking. These confounds make it difficult to draw conclusions about ability to suppress distractors based on speech-in-noise results. To more directly address the possibility of impaired distractor suppression, we applied a paradigm that provides behavioral and neurophysiological measures of auditory distraction to a cohort of tinnitus participants (Sorensen et al., 2025). This paradigm utilizes measures of neural synchronization to the external stimuli, which may be a particularly relevant feature to measure given the association between excess central gain and hypersynchrony (Hayes et al., 2021; McGill et al., 2022; Resnik & Polley, 2021).

## Methods

We utilized the STROBE cross-sectional guidelines in preparing this study for publication (von Elm et al., 2007).

### Participants

We recruited participants with tinnitus (n=38) to take part in the study. Participants were recruited via word of mouth, flyers, and advertisements on the Mass General Brigham Participant recruitment website. All procedures were approved by the Mass General Brigham Institutional Review Board and took place at Mass Eye and Ear between November 2023 and February 2025. After providing informed consent, study participants were screened for normal cognitive function (telephone Montreal Cognitive Assessment ≥18), English fluency (self-reported), mental health status (Beck’s Depression Inventory total score < 31), middle ear status (unremarkable otoscopy), and hearing status. Pure tone audiometric thresholds were required to be normal between 0.5 Hz and 2 kHz, corresponding to the range of our experimental stimuli, with no more than moderate hearing loss from 3-8 kHz. Those who were determined by an experienced clinician to have only intermittent tinnitus were excluded (n = 7), and one additional participant was excluded due to incorrect understanding of task instructions. The final count of 30 participants met our targeted recruitment based on a power analysis to detect a moderate effect between groups. Participants completed two sessions: an at-home testing portion completed on tablets (Microsoft Surface Pro 2, Pro 7, Redmond, WA) with custom testing software, followed by an in-person EEG session.

### Hearing Assessments

Participants underwent hearing assessments in both the at-home testing and in the lab under the direction of a licensed clinician. Assessments included the tinnitus handicap inventory (THI), hyperacusis questionnaire (HQ), pure tone audiometric threshold measurements, and measurements of loudness discomfort levels with the experimental stimuli to ensure that stimuli would not be presented at uncomfortable presentation levels.

Participants in this study were also asked about interest in participating in another study that did additional hearing assessments, including an assessment of time-compressed (65%) word recognition with reverberation (Noffsinger et al., 1994). For participants who chose to participate in this additional study (n = 22), we report these results.

### Experimental Stimuli

#### Target stimuli

Target stimuli consisted of concatenated AM tones with a constant carrier frequency and one of five AM rates (27, 41, 54, 68 or 81 Hz). Five AM tones, one at each rate, were ordered randomly to form a cycle. The same cycle was then repeatedly concatenated together to form a repeating pattern, or else different cycles were pseudorandomly concatenated (avoiding repeating the same AM tone back-to-back) to form random arrangements. Each of the 120 possible permutations of the five AM tokens was used as distinct pattern stimuli, with a corresponding number of random stimuli generated.

#### Melodic distractor stimuli

Melodic sequences were generated using the Magenta RL Tuner, a machine-learning based melody generator that incorporates explicit rewards for note sequences that follow Western musical theory in addition to patterns found in its training data (Jaques et al., 2017). Note sequences were transposed as pure tones in a region below the target stimuli, maintaining a 1/3 octave band below the lower sideband of the 81 Hz AM tone to minimize energetic masking in the frequency domain. This same sequence was copied 3 octaves higher to be in a region above the target stimuli, again maintaining a 1/3 octave band above the upper sideband of the 81 Hz AM tone.

Sequences that could not be transposed into this range were rejected. SAM was then applied to the tones. Note durations and SAM rates were adjusted between the at-home and in-person sessions to maintain an integer number of notes and AM cycles per note across the different hardware.

#### Noise distractor stimuli

Noise distractors were created to match the magnitude spectra of melodic distractors. From a given melody, the frequencies of the pure tones comprising the notes in the melody were taken as components of the noise distractor. Each component was present throughout the duration of the stimulus and the magnitude was scaled to the proportion of beats in the melody that contained the corresponding note. The starting phase of each component was randomized. Finally, the amplitude envelope from the melody, consisting of rest periods of zero amplitude and AM during note presentations, was applied to the noise stimulus.

### At-Home Procedures

In the at-home testing, participants were asked to report whether the target stimulus on each trial was randomly arranged or a repeating pattern in a speeded-choice task. Stimuli lasted 4 cycles (∼3 s) and were presented via calibrated Sennheiser HD280 (Wedemark, Germany) circumaural headphones. Responses were speeded by use of a scoring mechanic based on response time. At the start of each trial, a countdown started from 1000 and ran down until a response was recorded. Upon a correct response, the number remaining on the countdown was added to the total score. For an incorrect response, the number remaining on the countdown was subtracted from the total score. If participants failed to respond within 1.5 s of the end of stimulus presentation, 350 points were subtracted from the total score. The countdown was not visible during the trial but was shown with feedback after the response. Participants completed 8 blocks of 10 trials in each of the three conditions (target alone, with noise distractor at 12 dB SNR, and with melodic distractor at 12 dB SNR) in random order for a total of 240 trials.

Prior to beginning testing, participants went through several practice stages to familiarize themselves with each feature of the task. At each stage, participants needed to either respond correctly in 7 out of 10 practice trials, or complete 50 total practice trials to advance. In the first stage, participants were introduced to the target stimuli and practiced responding without any time pressure. In the second and third stage, participants were introduced to the two types of distractors and practiced responding to the target with the noise distractors and the melodic distractors, respectively. In the final stage, participants were introduced to the scoring mechanic that encouraged fast responses and practiced responding to the target stimuli alone with the scoring in place.

### In-Person EEG Session

The task was modified for the EEG session. Participants responded to target stimuli that either changed from random to repeating patterns after 4 cycles or stayed random throughout. Stimuli in the EEG session lasted a total of 12 cycles (∼9 s), and the polarity of the carrier frequency was inverted every cycle. Participants were instructed to refrain from responding until after the stimulation was completed, and the touchscreen participants used to record responses (Microsoft Surface Pro 4, Redmond, WA) did not display the digital buttons during auditory stimulation. Participants completed two blocks, one with the target presented with melodic distractors and one with the target presented with noise distractors, each at 12 dB SNR. Each block consisted of 240 trials, 120 trials each of random and random-repeating stimuli. EEG from a 64-channel BioSemi Active Two system (Wilmington, NC) was captured as participants completed the task in a sound-attenuated booth. Auditory stimuli were presented bilaterally through calibrated EARTone 3A insert ear headphones (Oaktree Products, Chesterfield, MO).

### EEG Pre-processing and Analysis

Recorded EEG data were imported into Matlab (Mathworks, Natick, MA) via EEGLAB (Delorme & Makeig, 2004). Data were filtered using fourth order Butterworth filters between 1-3000 Hz and rereferenced to the average of two mastoid electrodes. Epochs were selected for each level of analysis: cycle-length epochs for analyzing the frequency following response (FFR), single-token length epochs for analyzing the envelope following responses (EFRs), non-overlapping two-token length epochs for analyzing the envelope change following response (ECFR), and trial-length epochs for analyzing the distractor EFR and melody beat. For each level of analysis, we calculated the phase locking value (PLV) across epochs, taking the root mean square PLV across all channels. To compare across groups and conditions, the asymmetry index between the noise distractor and the melodic distractor conditions was calculated for each synchronization measure.

### Statistical Analysis

Comparisons between groups were done via two-sample t-tests, Mann-Whitney U tests, and repeated measures ANOVA, as appropriate. Statistical analyses were done in Matlab using the ttest2, ranksum, and ranova functions, with distractor type specified as the within-subjects factor and THI group as a between-subjects factor. The Mann-Whitney U test was used in lieu of two-sample t-tests when violations of normality were observed. We control the family-wise error rate using the Bonferroni-Holm method for repeated two sample t-tests making comparisons between the same conditions, although this correction did not change any significance test. To better fit the assumptions of parametric tests, rationalized arcsine transforms were applied to the behavioral accuracy data (Studebaker, 1985). Accuracies are plotted in the figures in rationalized arcsine units, but raw percentages are reported in the text for the ease of the reader.

## Results

### THI Groups

To assess how auditory distraction differs as a function of tinnitus burden, we split participants into two subgroups based on the median THI score within our sample (THI < 16, n = 16; THI > 16, n =14; Figure 1A). When referenced to the nominal THI categories, participants in the Low THI group correspond to a ‘slight’ tinnitus burden, while those in the High THI category would be categorized with a tinnitus burden ranging from ‘mild’ (18-36) to ‘severe’ (58-76). However, it is important to point out that these categorical boundaries and labels were only intended as rough approximations of tinnitus distress and are by no means deterministic (McCombe et al., 2001). Participant ages were matched between tinnitus severity groups, albeit with greater spread in the Low THI group (mean and standard deviation of Low THI group 53.8 ± 13.6 years; mean and standard deviation of High THI group 53.9 ± 8.3 years). Hearing thresholds did not vary between the Low THI and High THI groups (*F*(1, 28) = 0.36, *p* = 0.55; Figure 1B). The High THI group showed a greater tendency for reduced sound level tolerance based on the median HQ value (Low THI group median = 8; High THI group median = 10.5), though the difference was not statistically significant (Mann-Whitney U *z* = -1.36, *p* = 0.18; Figure 1C). Time-compressed word recognition scores from available subjects also did not differ between groups (*t*(20) = 0.1963; *p* = 0.25; Figure 1D).

**Figure 1:**
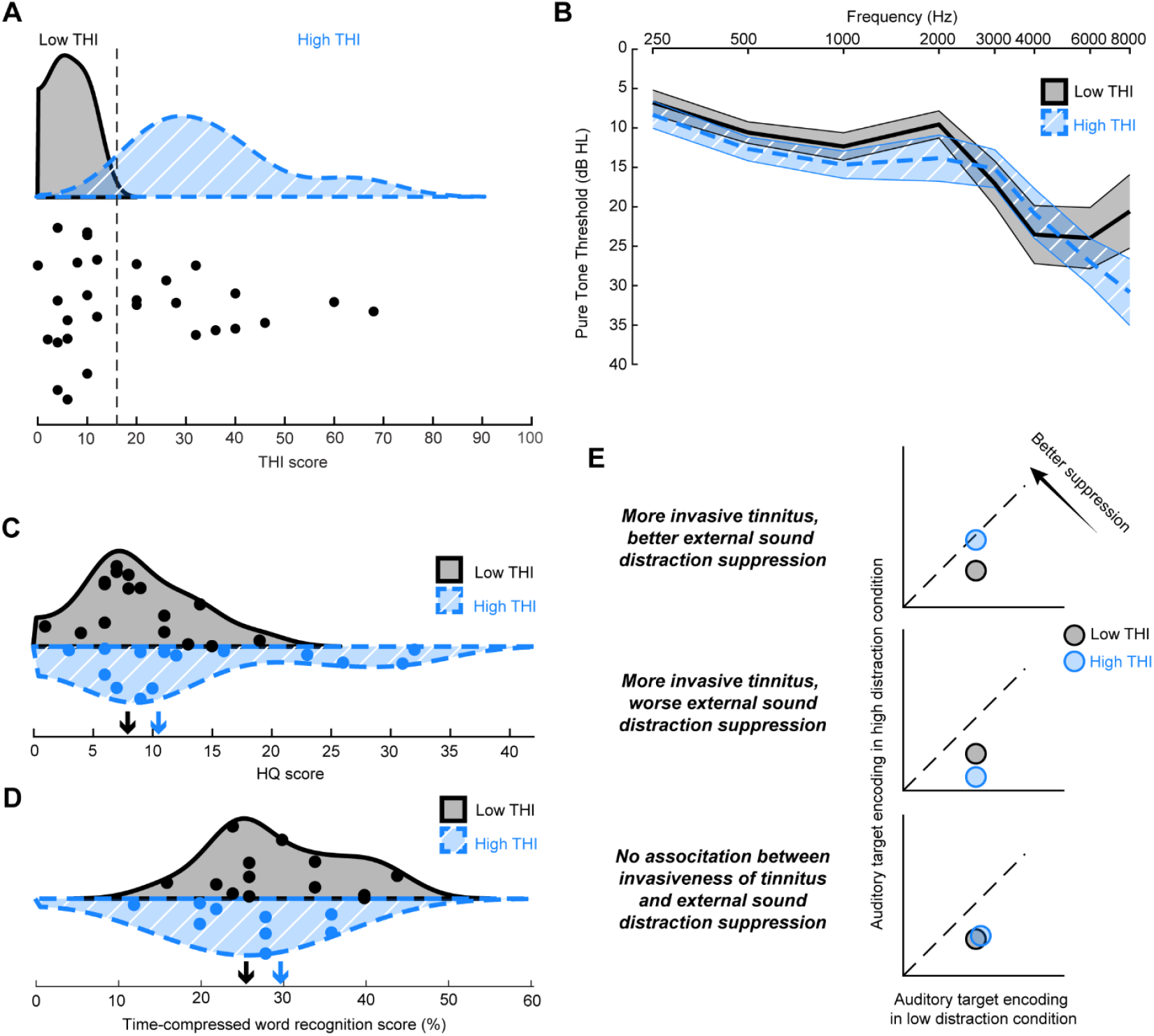
Variation in tinnitus burden is not readily explained by hearing status but may involve susceptibility to auditory distraction. **(A)** THI scores vary across our cohort of participants, with the median score below sixteen points (corresponding to a clinical grade of “slight or no handicap”). **(B)** Pure tone threshold audiograms are matched across the groups with lower and higher THI scores. Shaded regions depict the standard error of the mean. **(C)** While some individuals with high THI scores unsurprisingly also experience loudness hypersensitivity, the groups do not differ in median HQ scores. Arrows indicate group medians. **(D)** Word recognition scores from time compressed speech with added reverberation do not differ between the groups. Arrows indicate group means. **(E)** Model of how tinnitus may interact with distractor suppression. Top: Participants with more bothersome tinnitus may have extra practice suppressing distractors and have improved distractor suppression. Middle: Participants with more bothersome tinnitus may be unable to suppress distracting sounds and experience worse distractor suppression. Bottom: Suppression of an external sound may be unrelated to how the invasiveness of the internally generated phantom.

Thus, our participant cohort provided an opportunity to contrast differences in tinnitus severity while accounting for the potentially confounding influences of age, sound level tolerance, and hearing abilities. Theoretically, one could posit three relationships between the perceptual salience of the tinnitus phantom sound and distracting environmental sounds. The first possibility is that– on account of this indefatigable perceptual assault from their phantom sounds – persons with more bothersome tinnitus are better able to suppress more distracting environmental sounds (Figure 1E, top). The second possibility is that persons experiencing their tinnitus phantom as constant and irrepressible are less able to suppress more distracting environmental sounds (Figure 1E, middle). Yet a third possibility holds that the neural and perceptual salience of tinnitus phantom sounds are wholly separate from the processes that regulate environmental sounds, such that one will have no relationship with the other (Figure 1E, bottom).

### Behavioral Response to Distraction

To test these possibilities, we implemented a paradigm to measure the effects of differing levels of distraction (Figure 2). Participants listened to the target stimulus composed of a sequence of tones with different AM rates, and reported whether the AM tones are arranged into a random sequence or a repeating pattern (Figure 2A). The target stimulus was presented concurrently with an additional sound designed to impose variable degrees of distraction without introducing energetic masking: A melodic distractor and a noise distractor (Figure 2B and 2C, respectively). The melodic and noise distractors were matched in their magnitude spectra and thus provide the same long-term acoustic input, but our prior report demonstrated that the melodic distractor imposed a greater level of distraction, as evidenced by increased error rates and reaction time when neurotypical participants were asked to behaviorally report whether the target stimulus was arrayed in a random or patterned sequence (Sorensen et al., 2025).

**Figure 2:**
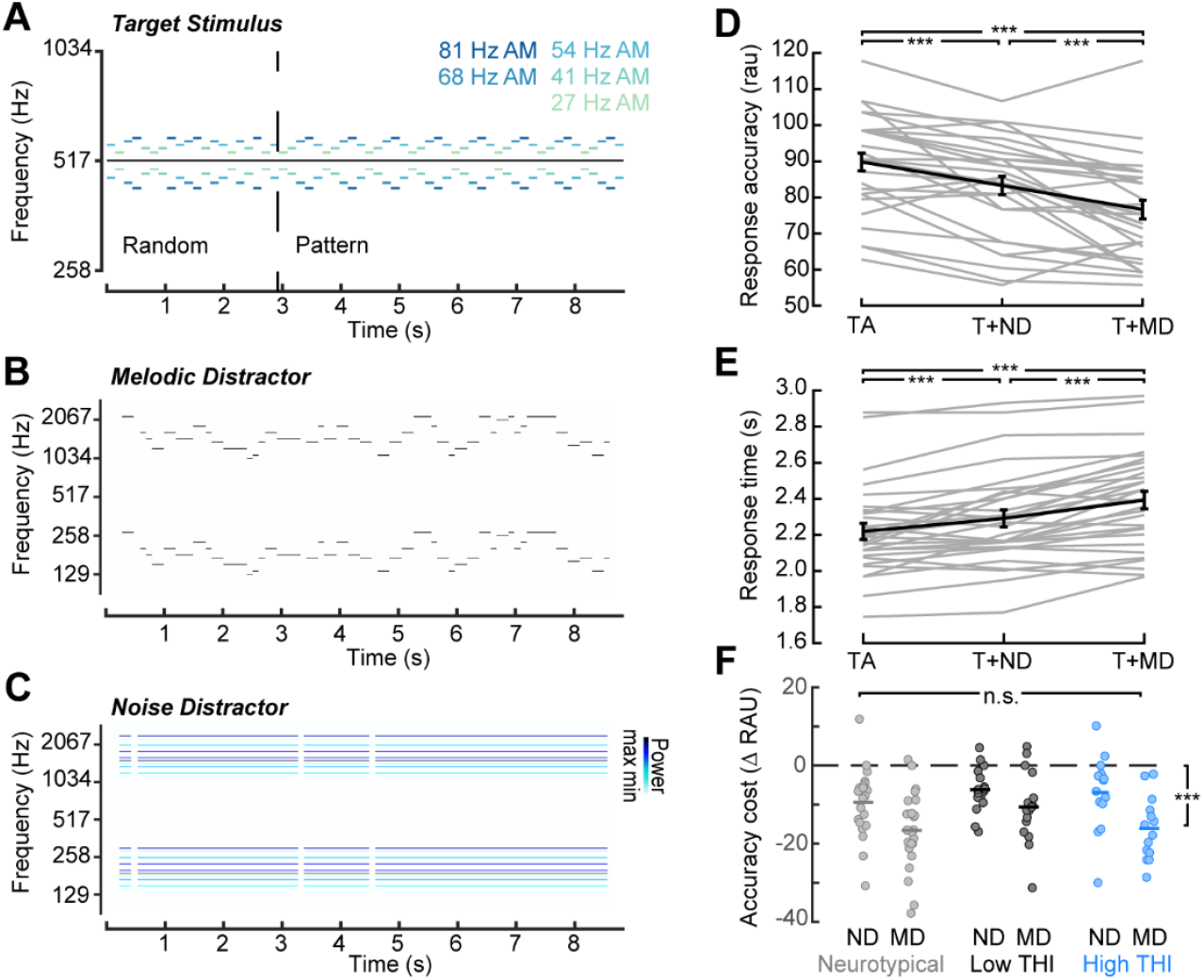
A novel paradigm to measure auditory distraction. **(A)** The target consists of a sequence of AM tones that are either randomly arranged or formed into repeating patterns; participants report what kind of arrangement they heard. **(B)** The melodic distractor consists of pure tone melodies generated via the Magenta RL Tuner and transposed above and below the target stimuli. **(C)** The noise distractor is made from the tones present in a melody spread across the stimulus duration. In the experiment, the melodic and noise distractors are also amplitude modulated; the spectrograms of the distractor stimuli are shown here without amplitude modulation for visual clarity. **(D-E)** In a speeded-choice task, participants correctly report whether the target was arranged randomly or in a repeating pattern when presented with the target alone (TA). Responses get less accurate (D) and slower (E) with increasing levels of distraction. **(F)** Participants perform no different on the task based on tinnitus group (neurotypical, low THI, high THI). Error bars depict the standard error of the mean. ND – noise distractor, MD – melodic distractor, *** - *p* < 0.001

Individuals with tinnitus were able to complete this task when the target was presented alone (TA) with a mean accuracy of 87% and responded after less than 2 repeats of the patterned cycle (mean response time 2.2 s; Figure 2D-E). When the target was paired with the noise distractor (T+ND), performance was impaired in both accuracy (mean 82%) and response time (mean 2.3 s). Performance decreased further still when the target was paired with the melodic distractor (T+MD, mean accuracy 76%; mean response time 2.4 s, Figure 2D-E).

Repeated measures ANOVA confirmed that both accuracy (*F*(2, 58) = 35.1, *p* < 0.001) and reaction times (*F*(2, 58) = 37.5, *p* < 0.001) differed between the conditions. Pairwise Tukey-HSD comparisons show that target alone accuracy is statistically greater than with the noise distractor (*p* < 0.001) and the melodic distractor (*p* < 0.001), and that accuracy with the noise distractor is statistically greater than with the melodic distractor (*p* = 0.001). Pairwise comparisons for response time show the same effects, with target alone responses recorded faster than with the noise distractor (*p* = 0.001) and melodic distractor (*p* < 0.001), and responses with the noise distractor faster than with the melodic distractor (*p* < 0.001).

We then assessed the performance cost of the noise and melodic distractor in Low- and High-THI subgroups alongside prior data in neurotypical participants (n = 23) by quantifying the change in accuracy relative to the TA condition (Figure 2F). Repeated measures ANOVA with group as a between-subjects factor showed a significant main effect of distractor type (F(1, 50) = 1.6), with the melodic distractor affecting performance more than the noise distractor. Neither the main effect of group (F(2, 50) = 1.66, *p* = 0.20) nor the interaction between group and condition (F(2, 50) = 1.91, *p* = 0.16) were significant. Performance of both THI groups was in line with performance of the normal hearing subjects in this difficult perceptual task.

### Neurophysiological Response to Distraction

While we did not observe any behavioral differences, there may be differences in the underlying neural representations that are compensated for behaviorally. To assess neural encoding, we measured EEG synchronization to nine different stimulus target and distractor features. Neural encoding of the target stimulus produced seven independent response features: The FFR to the constant carrier frequency of the target stimuli; EFRs to each of the five AM rates of the target stimuli; and an envelope change following response ECFR synchronized to the rate at which the target stimuli changes between different AM rates (Figure 3A-D). We were also able to independently and concurrently measure neural synchronization to the distractor stimuli; the noise and melodic distractors both produced an EFR to the AM rate at 19.5 Hz, and the melodic distractor also generated a following response at the beat rate (9.75 Hz). Our prior report had shown that of these nine features, the ECFR is sensitive to the level of distraction: it is decreased when paired with the melodic distractor compared to the noise distractor (Sorensen et al., 2025). We replicated this effect in tinnitus participants by observing that the neural target ECFR phase locking value was significantly more reduced during the melodic distractor than the noise distractor (paired t-test *t*(29) = - 4.53, *p* < 0.001; Figure 3E).

**Figure 3:**
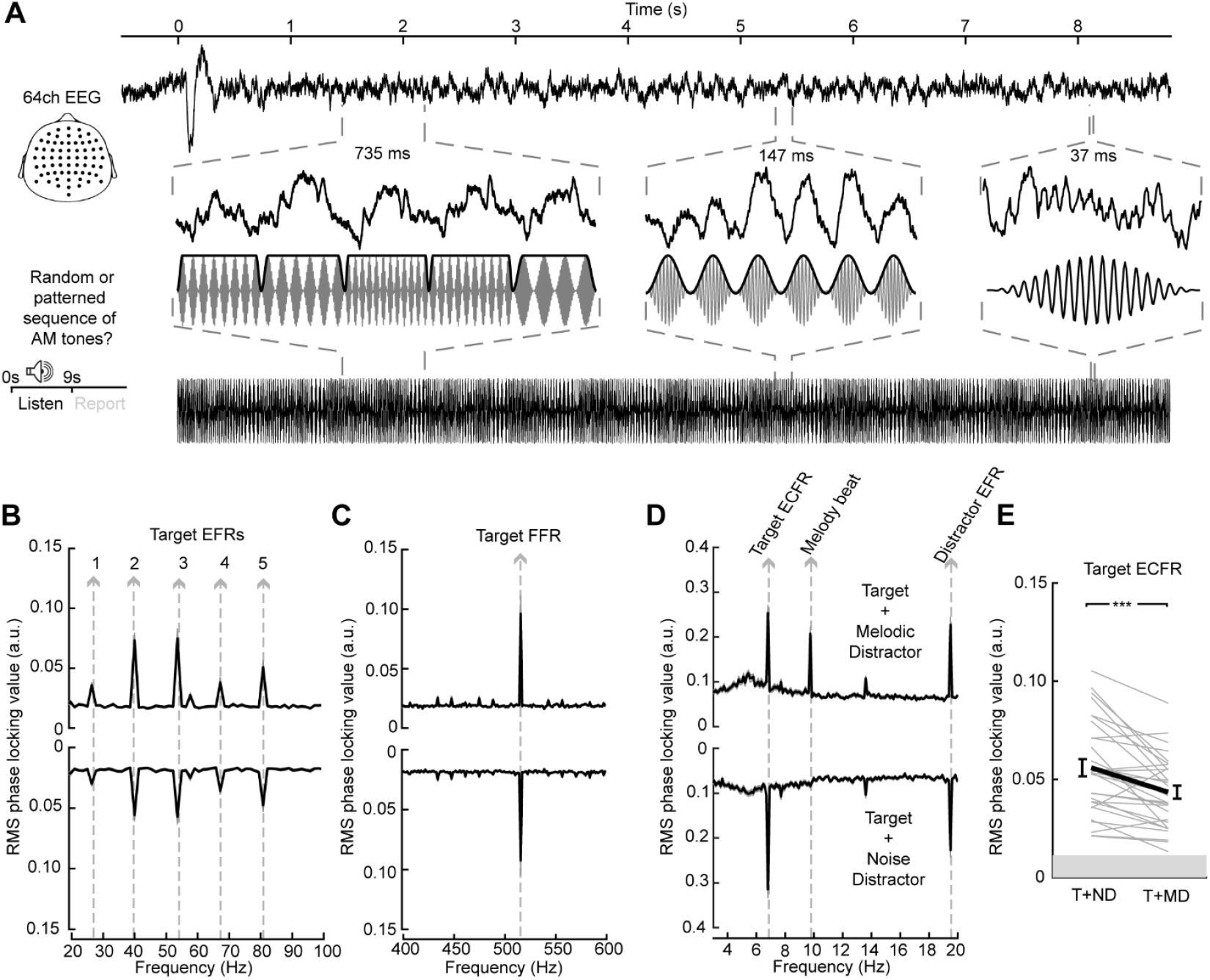
Measuring neurophysiological changes to distraction. **(A)** Synchronization in the EEG (top) to features of the target stimuli (bottom) are observed as participants perform the task. These include the ECFR, synchronized to changes between AM rates; EFR, synchronized to the amplitude modulation; and FFR synchronized to the carrier frequency. **(B-E)** Synchronization to target and distractor features can be quantified using the intertrial PLV. Synchronization measurements include EFRs to each of the five target AM rates (B); the target FFR (C); and the target ECFR, melody beat following response, and distractor EFR (D). Of these, the target ECFR is uniquely sensitive to the differing level of distraction between the melodic distractor and noise distractor (E). Error bars depict the standard error of the mean. *** - *p* < 0.001

### Responses to Distraction by Tinnitus Burden

Having established a neurophysiological paradigm to assess neural encoding of target sounds in the presence of competing sounds that provide variable levels of distraction, we next examined differences in perceptual and neural target salience during the active listening EEG session in Low THI and High THI participants. Behavioral distraction effects in the EEG session was similar to the effects observed during the speeded reaction time test (Figure 2F): both groups showed reduced accuracy with the melodic distractor (mean accuracy in low THI group = 84%; in high THI group = 87%) compared to the noise distractor (mean accuracy in low THI group = 93%; in high THI group = 95%; Figure 4A). Neither the interaction between group and distractor type (*F*(1, 28) = 0.176; *p* = 0.68) nor the main effect of group (*F*(1, 28) = 0.17; *p* = 0.68) were significant. The main effect of distractor type, however, was (*F*(1, 28) = 1013; *p* < 0.001).

**Figure 4:**
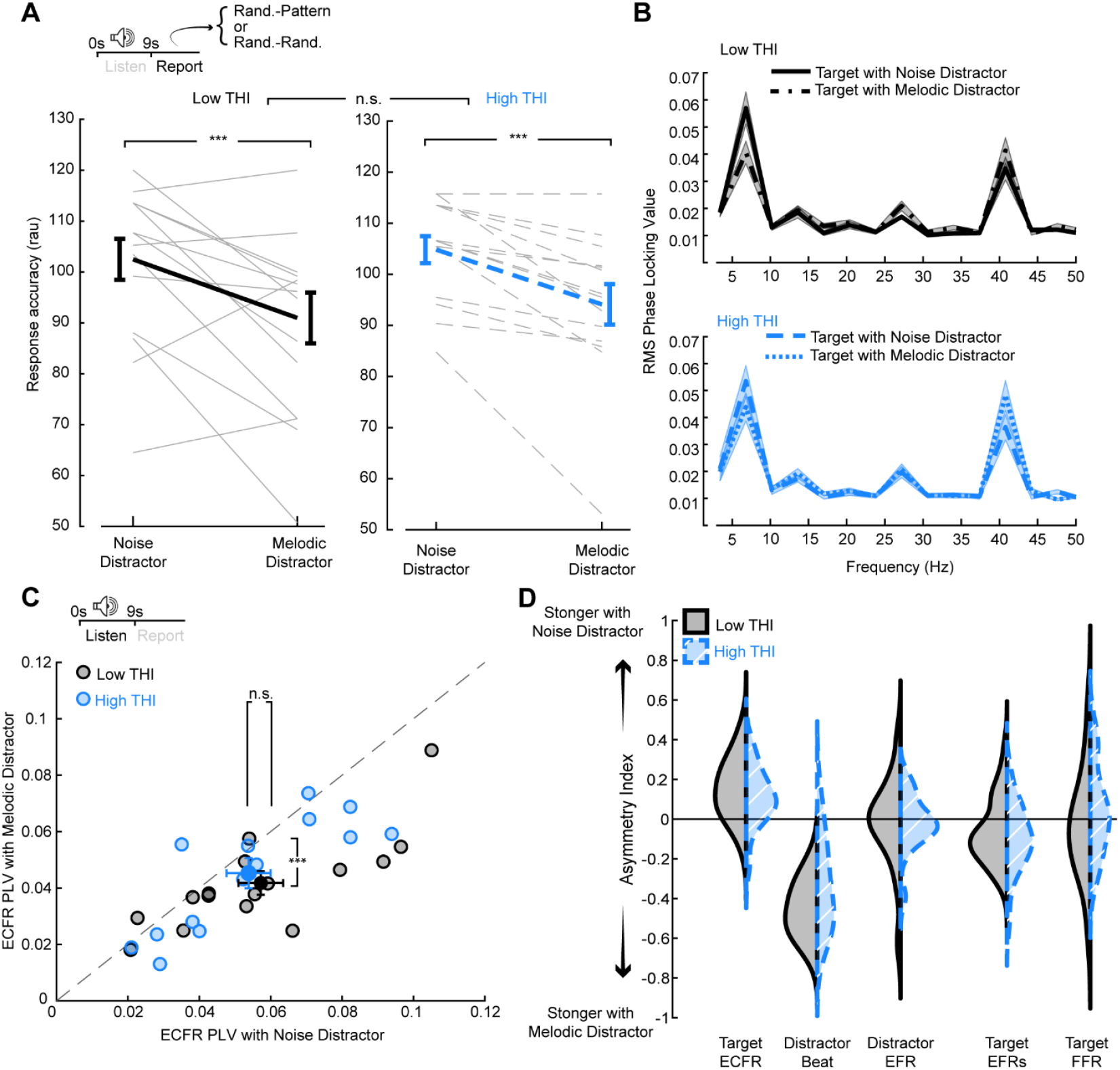
Participants with differing levels of tinnitus distress share common levels of behavioral and neurophysiological distraction. **(A)** While participants in both groups were less accurate at classifying the target stimuli when paired with the melodic distractor, no differences were observed between groups. **(B)** PLV spectra highlighting the ECFR peak at 6.8 Hz for the low THI group (upper panel) and high THI group (lower panel) with the noise distractor and the melodic distractor. **(C)** Both groups show a decrease in the ECFR with the melodic distractor compared to the noise distractor, and the group means are not significantly different. **(D)** Synchronization to all features of both the target and distractor stimuli, plotted as the asymmetry index between the response with the noise distractor and with the melodic distractor, do not differ between groups. Error bars depict the standard error of the mean. *** - *p* < 0.001; n.s. – not significant

To capture potential neural differences in distractor suppression between Low THI and High THI participants, we first investigated the ECFR, as it was sensitive to the level of external distraction in neurotypical participants (Sorensen et al., 2025) and also showed clear overall differences in the combined cohort of tinnitus participants (Figure 3E). PLV spectra of the ECFR are shown in Figure 4B. Quantification of the ECFR peak confirmed that both groups showed a decrease in the ECFR during the melodic distractor condition compared to the noise distractor condition, with the low group decreasing from a mean PLV of 0.058 to 0.042, and the high group decreasing from a mean PLV of 0.054 to 0.045 (Figure 4C). Repeated measures ANOVA showed a significant main effect of distractor type (*F*(1, 28) = 184; *p* < 0.001). However, the effect on the ECFR of increasing distraction did not differ between the low and high THI groups (group by distractor type interaction *F*(1, 28) = 0.03; *p* = 0.87), nor were the groups significantly different (main effect of group *F*(1,28) = 2.28x10-5; *p* = 0.99).

We next investigated neural synchronization to higher frequency modulations contained in the carrier frequency or amplitude modulation rate of the target and distractor stimuli (Figure 4D). The peak at the melody beat was only present in the melodic distractor condition, leading to a negative asymmetry index, but this did not differ between low and high THI groups (two-sample t-test *p*-adj. = 0.35). Asymmetry indices for the distractor EFR, target EFRs, and target FFR were near zero, indicating no effect of distractor type, and no differences were found between the groups for these measures (*p*-adj. = 0.50, 0.38, and 0.83, respectively).

## Discussion

Using a novel behavioral and neurophysiological paradigm aimed at measuring susceptibility to auditory distraction, we did not find evidence that individuals with more bothersome tinnitus were distracted by external stimuli different than those with less bothersome tinnitus. This lack of difference is consistent with a separation between perception and encoding for externally-presented stimuli and internally-generated sounds (Figure 1E, bottom). These results have implications for the development of objective biomarkers of tinnitus and our understanding of how the tinnitus percept interfaces with external auditory stimuli.

One important limitation of our study is the use of the overall THI score to separate the groups. While we chose this approach to try to approximate groups of individuals with benign and more bothersome tinnitus, the overall THI score is not designed for this purpose. Rough grading can be done based on the THI score, but evaluation of tinnitus would generally require a more holistic approach (McCombe et al., 2001). This approach thus collapses important features of tinnitus experience but was useful as a first pass approximation.

Our work sits alongside a number of other attempts to measure tinnitus via presentation of an external stimulus that fail to associate with tinnitus severity. For example, one might assume that the tinnitus is perceived as louder in persons with bothersome tinnitus, yet psychophysical assessments of tinnitus loudness have repeatedly shown no differences in tinnitus loudness after psychophysical matching nor any correlation with tinnitus severity indicated by questionnaire score (Martines et al., 2010; Smalt et al., 2022; Wallhäusser-Franke et al., 2012). Assessments of other tinnitus characteristics, such as minimum masking level and tinnitus pitch, also fail to associate with tinnitus distress (Karatas & Deniz, 2012; Meikle et al., 1984). One potential pitfall of assessments of the tinnitus percept itself is the inherent subjectivity and difficulty of matching an external sound to an internal sound. While such measurements can be stable over time (Fournier & Hébert, 2013; Nageris et al., 2010), there is no objective way to measure the “goodness of fit” to the tinnitus percept.

Perceptual responses to other types of external stimuli have also not shown strong associations with tinnitus. While a recent meta-analysis did suggest a slight decrease in speech-in-noise thresholds for subjects with tinnitus compared to normal hearing controls (Madhukesh et al., 2024), this has not been universally observed (Smith et al., 2025; Zeng et al., 2020). The estimated difference of about 3 dB SNR from the meta-analysis is of borderline significance, with larger differences usually required to motivate seeking treatment in individuals without tinnitus (McShefferty et al., 2015; Whitmer et al., 2016). In a comprehensive effort to measure differences in auditory perception in participants with tinnitus compared to normal hearing controls, Zeng et al. (2020) tested gap detection, frequency discrimination, intensity discrimination, temporal masking, amplitude modulation detection, and speech in noise recognition. Across all these measures, only frequency discrimination at 8 kHz was impaired in tinnitus participants, while intensity discrimination at 8 kHz, overshoot at 0 dB SPL/Hz noise, and speech perception in young participants with tinnitus were actually improved compared to control participants (Zeng et al., 2020). Another group showed that detection of a tone pitch-matched to the tinnitus frequency was not impaired in tinnitus participants. (Borst & Goedegebure, 2025). Our results represent another auditory task in which individuals with tinnitus perform as well as normal hearing subjects and further show that those with more tinnitus distress do not perform better or worse than those with less tinnitus distress.

These results suggest that there may be a fundamental difference between how patients with tinnitus perceive and react to the internally generated phantom percept and externally presented sounds, such that trying to study tinnitus via presenting sounds to the ear may not be effective. One exception may be sounds with negative valence: Previous work has shown that presenting emotionally evocative sounds does reveal changed pupil dilations and facial movements which correlate with tinnitus distress (Smith et al., 2025). This highlights the importance of limbic involvement in tinnitus, which has previously been observed (Rauschecker et al., 2010).

In contrast to studies on the perceptual consequences, neurophysiological data have been more fruitful in associating changes with the presence of tinnitus (see Husain & Khan, 2023 and Elgoyhen et al., 2015 for reviews). Changes in brain anatomy have been observed in MRI data, both in terms of gray matter volume (Makani et al., 2022) and anatomical connectivity (Benson et al., 2014; Crippa et al., 2010; Gunbey et al., 2017), although results are not always consistent between studies. Functional data in the resting state have shown alterations in default-mode, auditory, and attention networks (Kok et al., 2022; Schlee et al., 2009). Resting state data may be particularly fruitful for studying tinnitus, as participants with tinnitus will commonly be perceiving tinnitus at rest (although it may be masked by scanner noise in the case of fMRI). In quiet conditions using EEG, disruptions in some endogenous brain rhythms have been associated with tinnitus, particularly decreased alpha and increased gamma power in temporal cortex (van der Loo et al., 2009; Vanneste et al., 2010). Neural responses to externally-generated sounds have also shown some ability to differentiate tinnitus participants from normal hearing controls at early stages of the auditory pathway (Lanting et al., 2008; Melcher et al., 2000; Vasilkov et al., 2023), with manipulations of attention (Richardson et al., 2024), or in the processing of emotionally-evocative sounds (Carpenter-Thompson et al., 2014). While these other neurophysiological measures have shown associations with tinnitus, our neurophysiological measure of distraction employed in this study did not.

It should be noted that most studies compare individuals with tinnitus to individuals without, providing little insight into how any differences found may relate to tinnitus distress. While another attention-related biomarker of tinnitus in response to auditory stimulation has been reported (Richardson et al., 2024), no analysis was reported of the relation between the candidate biomarker and tinnitus distress. A marker that signifies tinnitus presence may be useful, but the best markers would be sensitive to tinnitus severity or distress. Some of the resting state EEG measures and fMRI measures in response to emotionally evocative sounds do associate with tinnitus severity (Carpenter-Thompson et al., 2015; van der Loo et al., 2009) and may be candidates to serve as a potential biomarker. Successful treatment of tinnitus is more often achieved by reducing distress experienced rather than eliminating the tinnitus percept altogether. While current assessments using self-report questionnaires are informative as to distress, self-reporting is subject to biases, leading to a particularly strong potential for placebo effects. Moving forward, biomarkers that focus on emotional and arousal responses to sound may prove a more fruitful avenue for sensitivity to tinnitus distress than the neutral stimuli utilized in this study and a number of past ERP studies. Interventions that aim to improve suppression of auditory distractors, such as those described by Searchfield et al. (2007) and Whitton et al. (2017), may also benefit from using distractors with negative valence rather than neutral distractors.

Our results indicate that individuals with tinnitus do not have a general deficit in suppressing auditory distractors. This suggests that there is something unique about the tinnitus percept that makes it difficult for those with greater distress to suppress. Limbic system involvement and negative valence associations may provide those unique qualities and are promising avenues to pursue for identifying ways to both measure and treat tinnitus distress.

## Acknowledgments

We acknowledge the input of Aravind Parthasarathy, Gerald Kidd, Hari Bharadwaj, Josh McDermott, and Barbara Shinn-Cunningham to the design of the paradigm used in this study.

